# Epstein-Barr virus nuclear antigen EBNA-LP is essential for transforming naive B cells, and facilitates recruitment of transcription factors to the viral genome

**DOI:** 10.1101/176099

**Authors:** Agnieszka Szymula, Richard D. Palermo, Ian J. Groves, Mohammed Ba abdullah, Beth S. Holder, Robert E. White

**Affiliations:** Section of Virology, Department of Medicine, Imperial College, London, London, United Kingdom.; Section of Pediatrics, Department of Medicine, Imperial College, London, London, United Kingdom.; Division of Infectious Diseases, Department of Medicine, Brigham and Women’s Hospital, Harvard Medical School, Boston, Massachusetts, United States of America.; Current address: Department of Pathology, Tennis Court Road, Cambridge, CB2 1QP, United Kingdom

## Abstract

The Epstein-Barr virus (EBV) nuclear antigen leader protein (EBNA-LP) is the first viral latency-associated protein produced after EBV infection of resting B cells. Its role in B cell transformation is poorly defined, but it is reported to enhance gene activation by the EBV protein EBNA2 in vitro.

We generated two sets of EBNA-LP knockout (LPKO) EBVs containing a STOP codon within each repeat unit of IR1. Intronic mutations in the first of these knockouts suggested a role for the EBV sisRNAs in transformation. LPKOs with intact introns established lymphoblastoid cell lines (LCLs) from adult B cells at reduced efficiency, but umbilical cord B cells, and naive (IgD+, CD27-) adult B cells consistently died approximately two weeks after infection with LPKO, failing to establish LCLs.

Quantitative PCR analysis of virus gene expression after infection identified both an altered ratio of the EBNA genes, and a dramatic reduction in transcript levels of both EBNA2-regulated virus genes (LMP1 and LMP2) and the EBNA2-independent EBER genes, particularly in the first 1-2 weeks. By 30 days post infection, these levels had equalised. In contrast, EBNA2-regulated host genes were induced efficiently by LPKO viruses. Chromatin immunoprecipitation revealed that recruitment of EBNA2 and the host factors EBF1 and RBPJ to all latency promoters tested was severely delayed, whereas these same factors were recruited efficiently to several host genes, some of which exhibited increased EBNA2 recruitment.

We conclude that EBNA-LP does not simply co-operate with EBNA2 in activating gene transcription, but rather facilitates the recruitment of several transcription factors to the viral genome, to enable transcription of virus latency genes. Additionally, our findings suggest that different properties of EBV may have differing importance in transforming different B cell subsets.

**Author summary:** Epstein-Barr virus (EBV) infects almost everyone. Once infected, people harbor the virus for life, shedding it in saliva. Infection of children is asymptomatic, but a first infection during adolescence or adulthood can cause glandular fever (mono). EBV is also implicated in several different cancers. EBV infection of B cells (the immune cell that produces antibodies) can drive them to replicate almost indefinitely (‘transformation’), generating cell lines. We have investigated the role of a virus protein – EBNA-LP – which is thought to support gene activation by the essential virus protein EBNA2.

We have made an EBV in which the EBNA-LP gene has been disrupted. This virus (LPKO) shows several properties. 1. It is reduced in its ability to transform adult cells, while immature B cells (more frequent in the young) die two weeks after LPKO infection. 2. Some virus genes fail to turn on immediately after LPKO infection. 3. Binding of EBNA2 to these genes is delayed, as is binding of some cellular factors. 4. EBNA-LP does not affect EBNA2-targeted cellular genes in the same way.

This shows that EBNA-LP is more important in immature cells, and that it regulates virus genes – but not host genes – more widely than simply through EBNA2.

## INTRODUCTION

Epstein-Barr virus is a ubiquitous human herpesvirus that asymptomatically infects the vast majority of the human population, particularly in the developing world, where primary infection typically occurs during the first few years of life, leading to lifelong EBV latency. Where primary infection is delayed into adolescence or adulthood, it can result in the temporarily debilitating but relatively benign condition, infectious mononucleosis. The major disease burden caused by EBV is the range of malignancies with which it has been associated. In particular EBV contributes to high levels of Burkitt lymphoma in sub-Saharan Africa and of nasopharyngeal carcinoma in southeast Asia, as well as around half of Hodgkin lymphoma cases, approximately one in ten gastric cancers, a range of B cell lymphomas in the immunosuppressed and more rarely with T and NK cell malignancies. Taken together EBV is implicated in around 1-1.5% of worldwide cancer incidence [1].

These diverse malignancies likely arise due to defects at different stages of the virus life cycle, or perhaps infection of cell types not involved in the virus’s natural life cycle [2]. The core of the EBV lifecycle occurs with the B cell compartment. EBV infection activates B cells, transforming them into proliferating lymphoblasts. In vitro these continue to proliferate into lymphoblastoid cell lines (LCLs) whereas in vivo they can differentiate - probably via a germinal center - into resting memory B cells where the virus is quiescent, producing RNAs but no viral proteins [3,4]. LCLs express the ‘growth’ program of EBV genes (latency state III), comprising six EBV nuclear antigens (EBNAs), the latency membrane proteins (LMP1, LMP2A and LMP2B) and a number of EBV encoded RNAs, including the abundant nuclear RNAs EBER1 and EBER2.

The latency III transcriptional program takes over 2 weeks to reach this state [5]. The first EBV proteins detectable after primary infection are EBNA2 and the EBNA leader protein (EBNA-LP) [6,7]. Shortly after this the EBNA3 proteins also become detectable [6], and the EBNAs rapidly reach the levels found in LCLs. In contrast, the LMP proteins take up to three weeks to reach LCL-like levels [5] and it has been proposed that the EBNA-normal/LMP-low transcription state that exists early after infection should be regarded as a new latency state - latency IIb [8]. EBNA transcription is initiated at multiple copies of Wp, the promoter in the major internal repeat (IR1) of EBV. Soon after infection, the burden of EBNA transcription shifts to Cp, a promoter upstream of IR1. Transcripts from both Cp and Wp are alternatively spliced, and translated in both cap- and IRES dependent manners to produce the six EBNAs.

The functions of most of the EBNAs are reasonably well understood: EBNA1 is important for the replication and segregation of the viral genome during the cell cycle, by binding to oriP. The EBNA1/oriP complex is also important in the switch from Wp to Cp [9]. EBNA2 is essential for the initial transformation of B cells, rapidly activating both host and viral genes through recruitment to promoters or enhancers alongside cellular transcription factors such as Pu.1 [10], RBPJ (also called CBF1) [11],[12], IRF4 and EBF1 [13,14]. The EBNA3 proteins are slow-acting transcriptional repressors that are important for suppressing senescence and apoptosis around 3 weeks after infection [15,16]. Notably, the EBNA3s and EBNA2 appear to have a close relationship with each other, co-regulating genes and being bound at the same chromosomal location [13,17-19].

In contrast to the other EBNAs, however, the role played by EBNA-LP in B cell transformation is not known. Through initiation at different Wp promoters and exon skipping in Cp transcripts, the EBNA-LP protein comprises a variable copy number of a 66 amino-acid N terminal repeat domain (encoded by exons W1 and W2 within IR1) and a C terminal domain encoded by exons Y1 and Y2. In LCLs, EBNA-LP mainly localizes to PML nuclear bodies (ND10) [20] although it takes several days after infection to accumulate there [21]. Functionally, EBNA-LP has been shown to enhance the activation of host and viral genes by EBNA2 after transfection, although not all studies agree on which genes are affected [7,22-26].

The complex repetitive nature of the EBNA-LP gene makes its analysis in the viral context challenging. Previous genetic analyses of EBNA-LP have been restricted to mutation of the C-terminal Y exons [27,28]. These Y domain knockout viruses establish LCLs at a much reduced efficiency, and then only when the early outgrowth of the cell lines was supported by growth on irradiated fibroblast feeder cells. Deleting increasing numbers of IR1 repeat units below five progressively reduced transformation efficiency [29], but as well as changes to maximum EBNA-LP size, this decrease could be due to the reduced Wp number producing less of the EBNA proteins (particularly EBNA2) or of the recently identified stable intronic RNAs (sisRNA1 and sisRNA2) that are produced from the introns between W exons [30].

These prior studies of EBNA-LP function have been conducted in the context of transfecting isolated genes, and/or in the presence of the truncated EBNA-LP protein produced by the P3HR1 virus, and not in the context of virus infection. Therefore, the aim of this project was to produce a complete EBNA-LP knockout virus, and use it to establish the importance of (and a role for) EBNA-LP in the transformation of B cells. While our first EBNA-LP knockout was additionally defective due to mutations in the introns between the EBNA-LP exons, a second, cleaner knockout showed that EBNA-LP is important but dispensable for the transformation of adult memory B cells, but is essential for the transformation of naïve B cells. Furthermore, both knockouts demonstrated that EBNA-LP is crucial for establishing and stabilizing the viral transcription program after infection, probably through facilitating the recruitment of EBNA2 and the host protein EBF1 to the incoming virus genome. However, EBNA-LP did not enhance the induction of host genes by EBNA2 during infection.

## RESULTS

### Generation and validation of EBNA2- and EBNA-LP-deficient BACs

Because of the multiple copies of Wp, and the alternative splicing of the EBNA-LP transcript, the only valid approach to completely knockout EBNA-LP was to introduce a nonsense mutation into the EBNA-LP coding region – with a PvuI restriction site to help screening – (Fig 1A) into each of the IR1 repeat units of EBV. This was done initially using class IIS restriction enzymes to generate an array of 6.6 mutated IR1 repeat units (Fig S1), the same size as in the parental EBV BAC (designated wild-type (WT)-HB9), and matching the typical size of IR1 in circulating viruses [31]. This approach necessitated the point mutation of a BsmBI restriction site in the short intron between exons W1 and W2 (Fig 1A). This mutant IR1 repeat was introduced into the viral genome using RecA-based recombineering, first deleting IR1 from WT-HB9 and then introducing the mutant repeat array into the viral genome. This produced the EBNA-LP-knockout virus LPKO^i^, where ‘i’ denotes the intronic point mutation of the BsmBI restriction site. A revertant virus (LPrev^i^) was made using a repeat array containing a wild-type W1 exons (i.e. encoding an intact EBNA-LP), but retaining the intronic point mutation, to control for any impact of this mutation. Two knockouts and their revertants were generated independently, as summarized in the flow chart (Fig S1).

**Fig 1.**
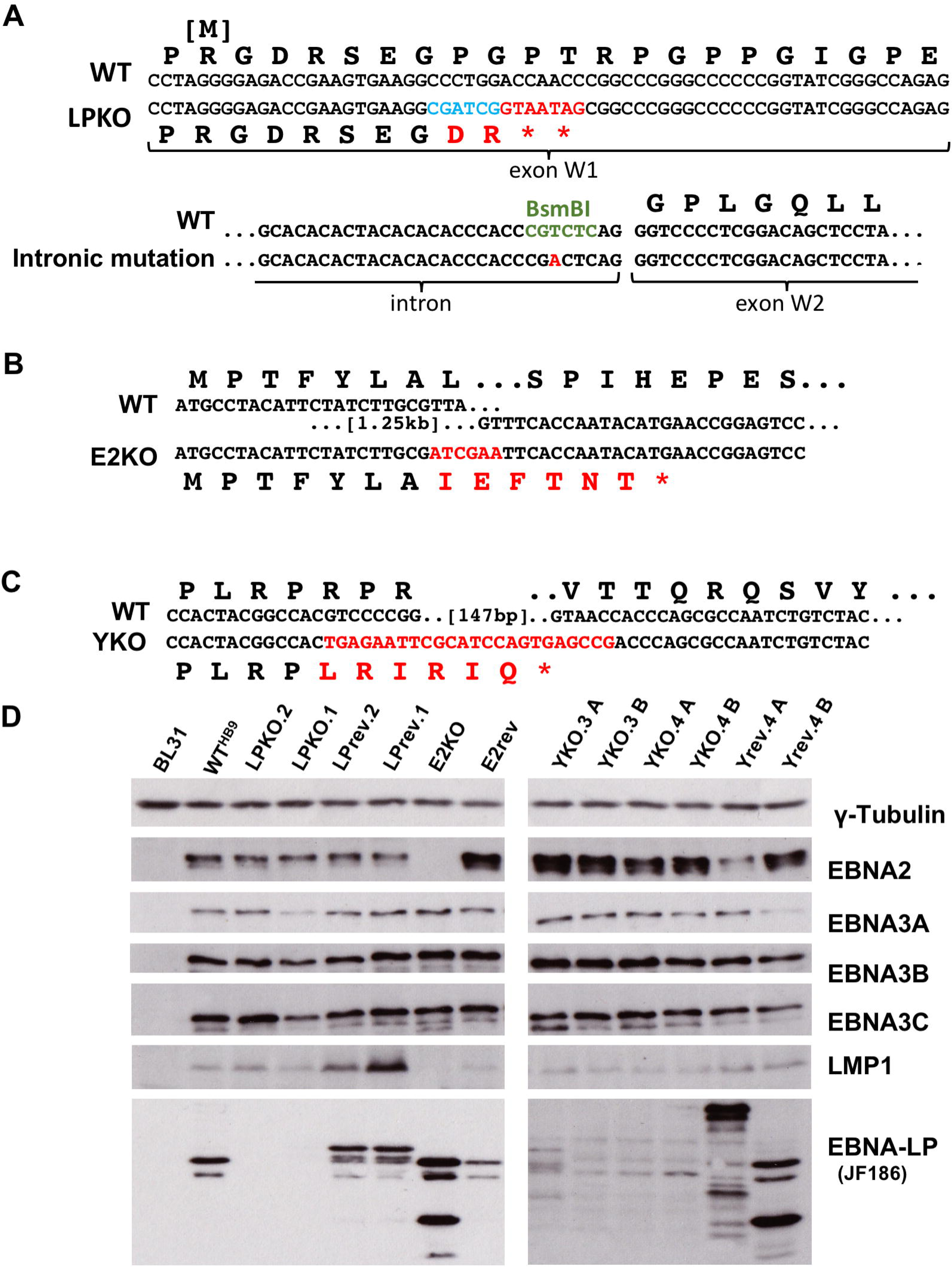
Construction and validation of EBNA-LP knockouts and their revertants. Sequence changes introduced in the production of: **A.** EBNA-LP knockout, LPKO^i^, and the intronic mutation also shared by LPrev^i^; **B.** EBNA2 knockout, E2KO; and **C.** the EBNA-LP truncation mutant YKO. Protein translations are shown above the nucleotide sequence, with the initiating methionine of EBNA-LP created by alternative splicing is shown in square brackets. The nucleotide changes (red) and the introduced PvuI restriction enzyme site (blue) are indicated. The BsmBI restriction site (green) deleted by a single T to A nucleotide change in LPKO^i^ and LPrev^i^ is indicated. **D.** Western blotting of EBV protein levels in BL31 cells stably infected with the various recombinant viruses. A and B suffixes indicate independent BL31 cell lines produced from the same virus.

In order to facilitate comparison with the previous genetic studies of EBNA-LP function in a P3HR1 strain backbone [27,28], we also generated a pair of recombinant viruses (designated YKO) that lacked the protein domains of the Y exons but retained exon Y1 splice acceptor and exon Y2 splice donor (Fig 1B). A revertant (Yrev) was generated for one of these knockouts. In order to separate the role of EBNA2 from that of EBNA-LP, an EBNA2 knockout (E2KO) EBV – and its revertant, E2rev – were also generated. E2KO retains the entire Y3 exon and its 3’ splice site. This allows qPCR detection of Y2-YH EBNA2 transcripts in the E2KO, despite being deleted for the rest of the EBNA2 ORF (Fig 1C). All of the BACs were screened by restriction digestion and pulsed field gel electrophoresis to ensure they were identical to WT-HB9 except for the intended modifications (Fig S2).

Infectious virus was produced from cell clones produced by transfection of recombinant BACs into 293 cells. The Burkitt lymphoma cell line BL31 – which we and others have previously used to establish cell lines for recombinant EBVs that are deficient in transformation [32,33] – was used to establish cell lines after infection with each virus. These cell lines did not apparently differ in the splicing of transcripts initiated at either Cp (Fig S3) or Wp (not shown), other than the expected shortening of transcripts in YKO cell lines caused by the deletion in the Y1 and Y2 exons. Similarly, the mutations did not alter the levels of any latency proteins other than those that had been mutated (Fig 1C and Fig S4). However, the YKO genomes only produced a very low level of C-terminally truncated EBNA-LP, and neither proteasome inhibition (MG132 treatment), nor analyzing whole cell lysates changed this observation (data not shown). We also noted a propensity for LPrev^i^ to produce larger sized and more abundant EBNA-LP isoforms, and that our EBNA2 knockouts exhibit higher EBNA-LP protein levels (Fig S4). However, overall it appears that in established BL31 cell lines, knockout of EBNA-LP does not alter the protein levels of other EBV latency genes.

### LPKO^i^ is unable to transform B cells into LCLs

Next, the ability of the recombinants to transform adult B cells was assessed. Under the microscope, uninfected B cells were indistinguishable from those infected with E2KO (EBNA2 knockouts are known to be completely transformation defective [34]), whereas LPKO^i^, YKO and revertants all induced an apparent activation of the B cells within 3 days of infection, characterized by cell enlargement and aggregation into clumps (Fig 2A). However, these aggregated cell clumps expanded over the next few days in the wild-type-infected cells whereas LPKO^i^ and YKO cell lines lagged behind in outgrowth (Fig 2B). We also noted that the expansion of both of the LPrev^i^-infected cells also lagged somewhat behind the other wild-type infections. Thereafter, YKO, LPrev^i^ and the other wild-type and revertant viruses were all reproducibly able to establish LCLs.

**Fig 2.**
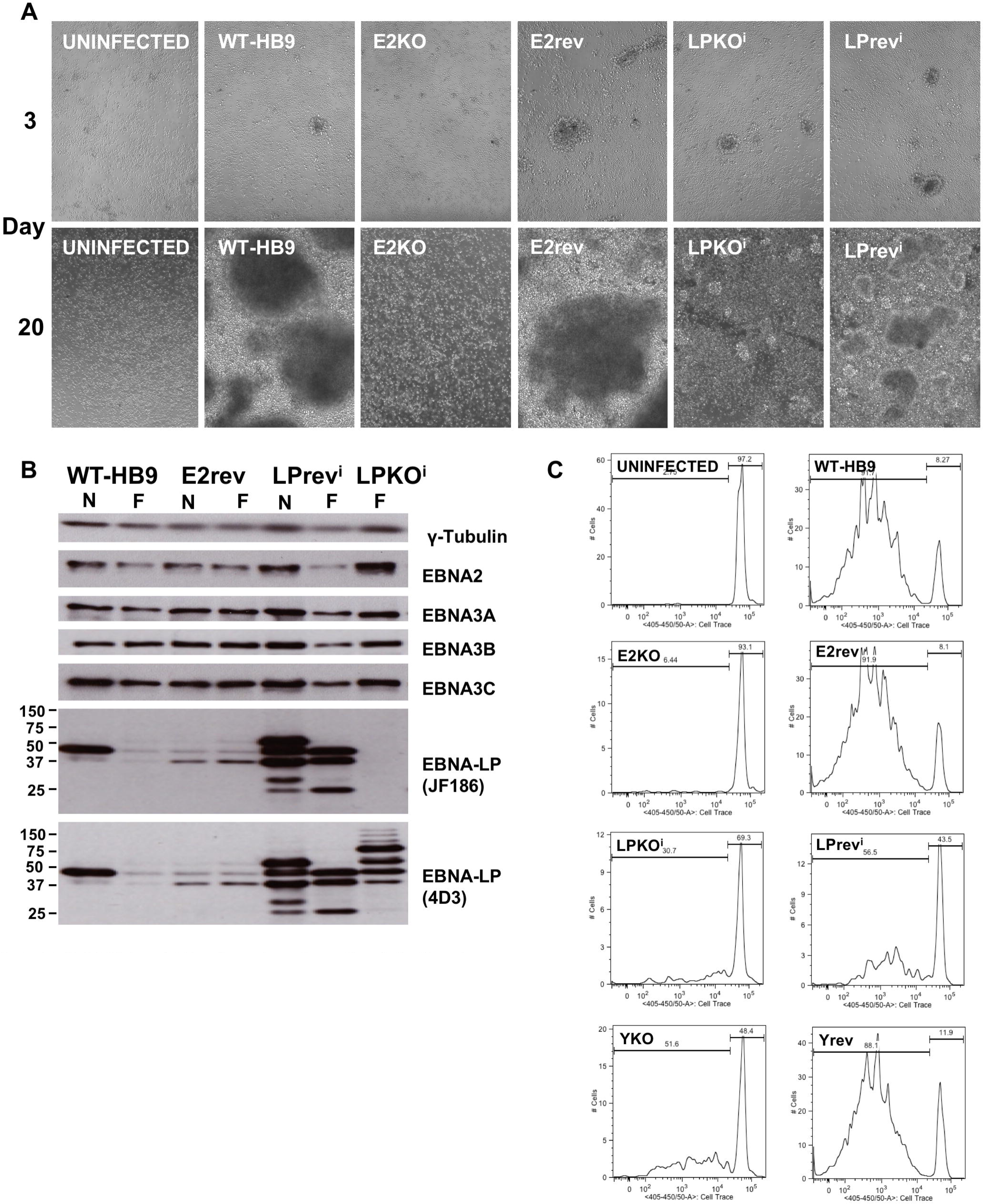
Both LPKO^i^ and LPrev^i^ are transformation-defective. **A.** CD19-selected adult B cells were infected with viruses as indicated. Cell activation and transformation as seen under 10x magnification at the time points indicated (see Fig S5 for more time points). **B.** Western blots of proteins from LCLs grown out from recombinant EBV infections. The virus used for the outgrowth is indicated. Initial phase of the outgrowth of cells was either performed on irradiated MRC5 feeder cells (F) or without feeder cells (N). The epitope in EBNA-LP recognised by the JF186 antibody exists in B95-8 but is missing from most virus strains. Antibody 4D3 recognises all known EBNA-LP variants. **C.** Flow cytometry plots from live CD20-positive cells 7 days after infection of adult B cells stained with CellTrace violet prior to infection. Degree of dilution of the violet signal is indicated on the x-axis, indicating number of cell divisions.

Western blotting of these LCLs confirmed the observation made in BL31 (Fig 1D) that YKO made only small amounts of truncated EBNA-LP. In contrast, LCLs were only established twice (in over 30 experiments) after infection with LPKO^i^. One of these was a spontaneous LCL that lost the LPKO^i^ genome during culture. The other was probably a coinfection between a (presumably) donor-derived virus and LPKO^i^, since it produced a variant of EBNA-LP that was not from B95-8 (Fig 2B), as well as LPKO^i^-derived transcripts (identified by cloning and sequencing), and the LPKO^i^ genome rescued from the cells into bacteria was identical to the parental BAC (not shown). Overall, this suggests that LPKO^i^ is unable to transform B cells into LCLs.

### LPKO^i^ supports limited proliferation after infection of naive B cells

In order to better understand the difference in transformation between the wild-type, LPKO^i^ and E2KO viruses, cell behaviour during the early times after infection was investigated further. Cell proliferation was tracked by measuring dilution of CellTrace^TM^ Violet over the first 10 days post infection. As previously reported [35] EBV-infected cells did not divide until after day 3 post infection (Fig S6), during which time cells in the LPKO^i^, revertant and wild-type infections increased in size and clustered together, whereas uninfected and E2KO cells remained largely unchanged (data not shown). From day 5 to 10 post-infection, increasing numbers of proliferated cells were seen in wild-type and revertant infections (Fig 2C and Fig S6). In contrast, only a few proliferated cells are apparent in the LPKO^i^-infected populations, while the YKO and LPrev^i^ viruses induced more proliferation than LPKO^i^ but less than the wild-type controls. These observations were consistent for both LPKO^i^/LPrev^i^ pairs.

This suggests that while EBNA-LP contributes to transformation, it is not required for many of the activation functions fulfilled by EBNA2, as the E2KO-infected cells were apparently as inert as uninfected cells. However, there is also some defect in the LPrev^i^ viruses that may also compromise the function of LPKO^i^. This might have been due to the intronic mutation of the BsmBI restriction site, but resequencing the repeat unit used to generate LPrev^i^ and LPKO^i^ revealed that there were three additional non-consensus nucleotides in the repeat. Analysis of B95-8 genome sequence has now demonstrated that these changes are found in a single repeat unit within the IR1 of both the WT-HB9 BAC and the original B95-8 cell line (Ba abdullah et al; manuscript submitted for publication). Furthermore, this one repeat unit also contains an EBNA-LP STOP codon at the end of exon W1 (Fig S7). Together, this means that none of the viruses described so far (including the widely used B95-8 BAC) have a truly intact IR1: WT-HB9 (plus E2KO, E2rev, YKO and Yrev) contain 5 intact repeat units, and one with a defective EBNA-LP exon pair and three non-consensus bases in BWRF1; LPrev^i^ contains six intact EBNA-LP exon pairs, but each repeat also contains four intronic mutations (one removing BsmBI in the small intron, and the three in BWRF1); in LPKOi, each repeat unit carries these intronic mutations and the stop codons designed into EBNA-LP.

### LCLs can be established using an improved LPKO-mutant EBV and its wild-type counterpart

In order to correct for the intronic defect of LPrev^i^, and assess whether it also altered the behaviour of LPKO^i^, we isolated an IR1 repeat unit that matched the B95-8 consensus sequence, and used it to generate two new repeat arrays – one wild-type and a second consisting the LPKO mutation described in Fig 1A – using a method based on Gibson assembly [36] that avoided mutation of the BsmBI restriction site in the small intron (Fig S8A): all of the IR1 sequence (other than the defined EBNA-LP mutations) matched the published B95-8 sequence. Both of these repeat arrays were recombined into the IR1-knock-out that had been used to generate LPKO^i^.2 to make two independent LPKO^w^ BACs (where ‘w’ indicates **w**ild-type IR1 backbone) and one with a wild-type repeat with no heterogeneity (WT^w^) (Fig S1C). These were validated by pulsed field gel electrophoresis (Fig S8B) and used to generate virus-producing cell lines.

LPKO^w^ and WT^w^ were used to infect adult B cells and compared with the previous viruses (Fig 3A and supporting Fig S9). At 8 days post infection it is clear that WT^w^ matches (and perhaps exceeds) the transforming capability of the parental wild-type BAC and revertants, and is considerably superior to LPrev^i^. More interestingly, LPKO^w^ is superior to LPKO^i^ in driving infected B cells to undergo proliferation, approaching the level seen for YKO, suggesting that many of the important functions lost in LPKO^w^ are also missing in the YKO infection.

**Fig 3.**
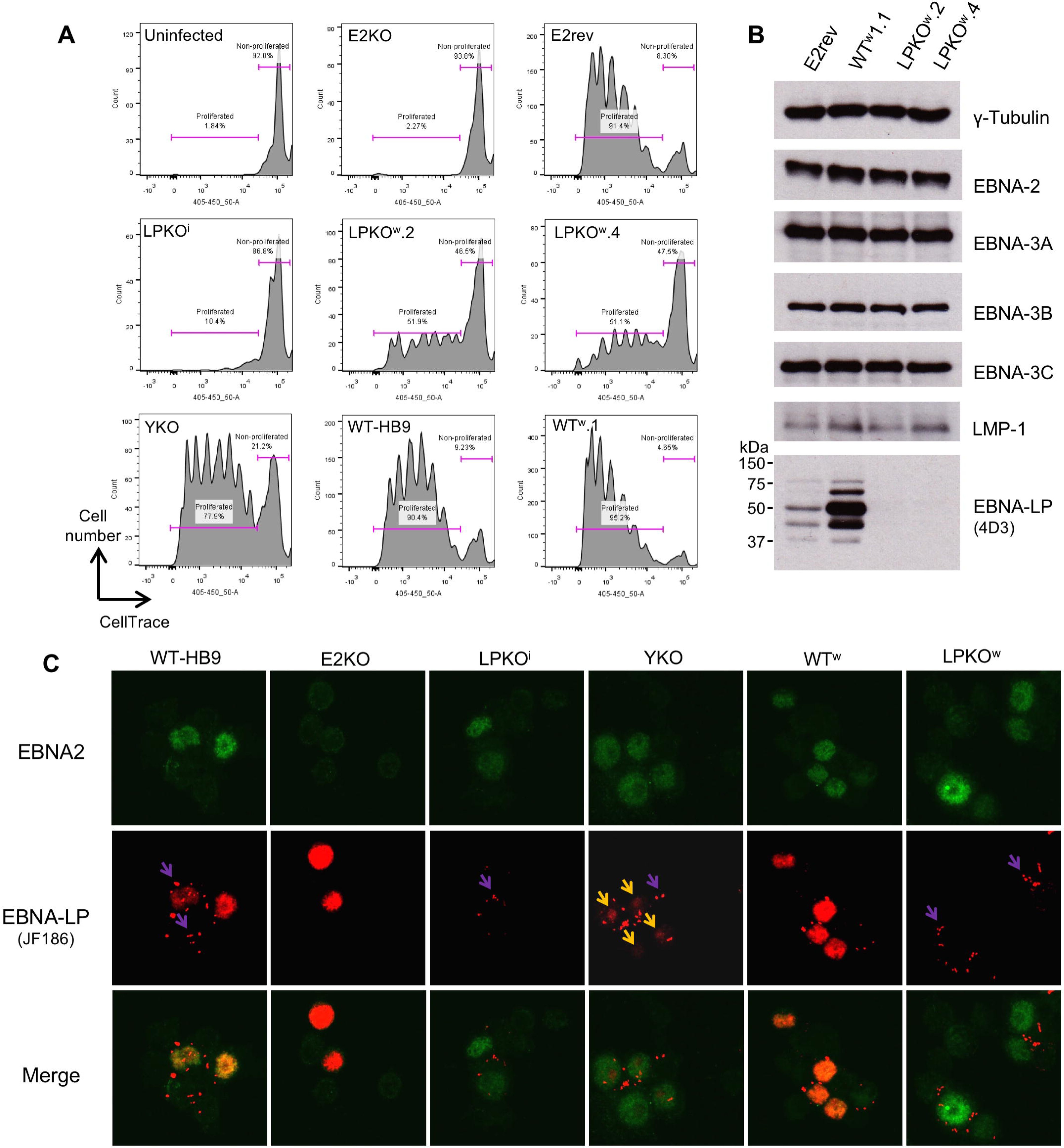
Both LPKO^w^ and WT^w^ are superior in transformation than LPKO^i^ and WT BAC respectively. **A.** Cell proliferation of live B cells 8 days post-infection, assessed by dilution of cell trace violet. **B.** Western blotting of viral proteins in LCLs established with LPKO^w^ and WT^w^ viruses. **C.** Immunofluorescence analysis of EBNA2 and EBNA-LP expression 48 hours post infection. Purple arrows indicate extracellular (or pericellular) foci that are artefacts also seen with the secondary antibody alone. Yellow arrows indicate nucleolar accumulation of EBNA-LP in YKO infections. The red single channel image in YKO has been brightened to improve visualisation of the faint nucleolar EBNA-LP signal. Other channels use the same brightness across the experiment.

Infection of 10^6^ B cells with LPKO^w^ at an MOI of 1 rgu/cell consistently induced expansion for approximately 5-7 days, but then appeared to stagnate for the next 1-2 weeks, after which cells usually proliferated again, and subsequently established LCLs. The other viruses with reduced transformation efficiency - YKO and LPrev^i^ – did not exhibit this period of lag in growth, and generally established LCLs more quickly than LPKO^w^. Latency protein levels were largely similar between the LCLs (Fig 3B), with LPKO^w^ LCLs clearly lacking EBNA-LP, and WT^w^ showing an elevated level of EBNA-LP relative to the parental wild-type.

The viruses were further validated by immunofluorescence analysis of B cells infected 48 hours post infection (Fig 3C), although this analysis was complicated by extra-cellular artefacts detected by anti-mouse Ig secondary antibodies. EBNA2 levels were similar in infected cells between all of the recombinant viruses (except E2KO, which – as expected – lacked EBNA2). In contrast, EBNA-LP levels were dramatically higher in E2KO-infected cells compared to wild-type infections, while the YKO EBNA-LP was expressed at much lower levels - consistent with western blotting of YKO LCLs and BL31 cell lines (Fig 2D) - but also appeared to be exclusively nucleolar (Fig 3C). EBNA2 protein levels in LPKO^i^- and LPKO^w^-infected B cells were indistinguishable by immunofluorescence (not shown).

### EBNA-LP mutant EBVs are defective in transforming cord blood and naive adult B cells

We also infected mononuclear cells from umbilical cord blood to try to establish LCLs. However, we were repeatedly unable to establish cord blood LCLs with either LPKO^w^ virus or the YKO virus, whereas WT-HB9, WT^w^ and the more defective LPrev^i^ all established LCLs consistently. The same effect was observed for infection of CD19-selected B cells from cord blood. In adult lymphocytes, LPKO^w^-infection resulted in more dying cells (i.e. sub-G1 DNA content) than WT^w^ infection, but in cord blood there were both more dead cells and – by day 11 – far fewer cells in S or G2 phases of the cell cycle (Fig 4A). By approximately 14 days post infection (precise timing varied with each donor), just as LPKO^w^-infected adult cells recommence their expansion, there appear to be no remaining live cells, as any remaining clumps of cells disintegrated and never recovered. This shows that cord cells arrest and die around 1-2 weeks after infection with an EBNA-LP-deficient EBV.

**Fig 4.**
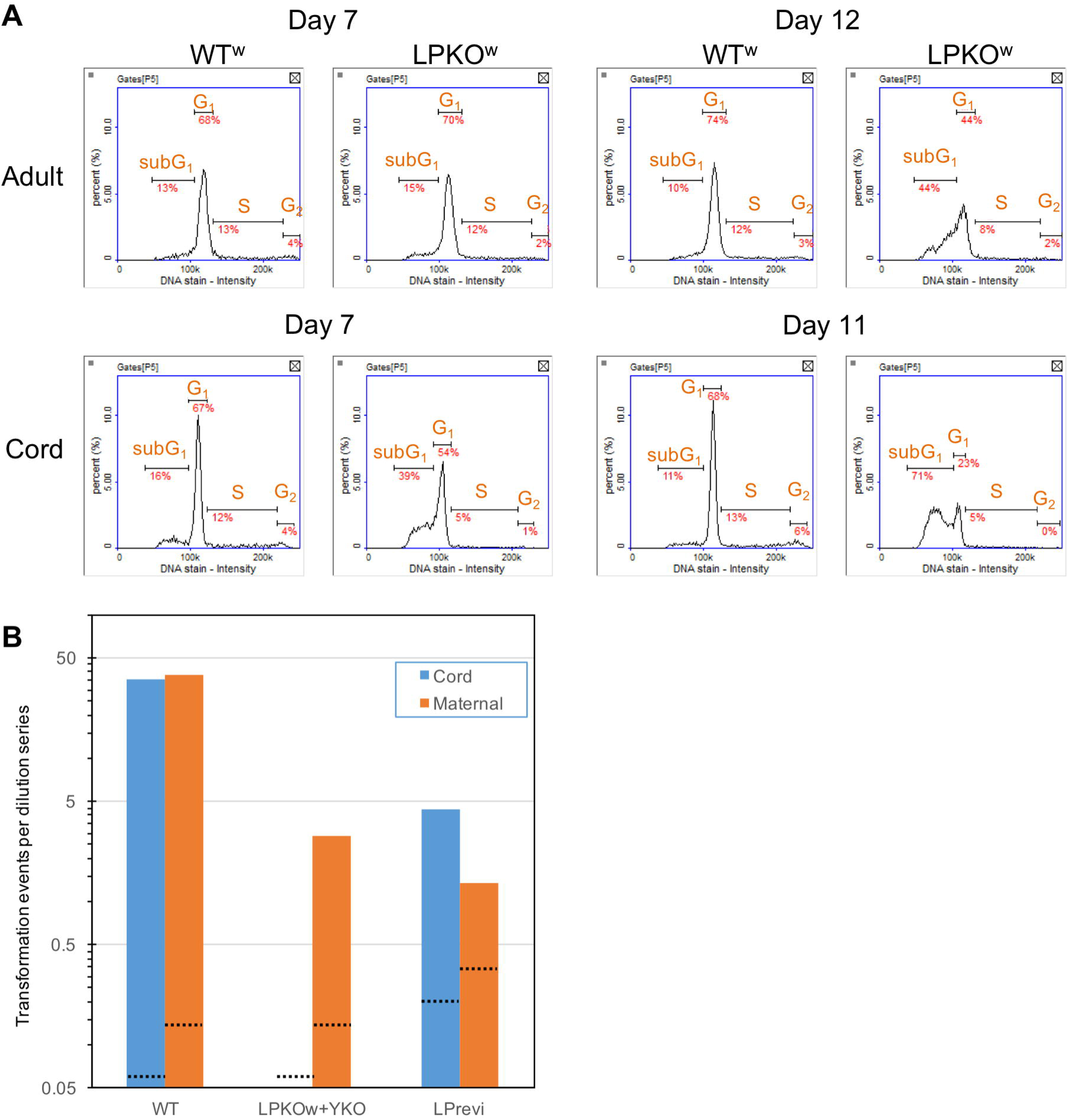
EBNA-LP mutants are defective at transforming B cells from cord blood. **A.** Cell cycle profiles of CD19+sorted adult and cord cells infected with WT^w^ or LPKO^w^ viruses. Graph shows the DNA quantity per cell (from DAPI staining) **B.** Transformation efficiencies for each infection were calculated from two-fold dilutions of infected cells (see Methods). These efficiencies were averaged for each virus group in each cell type across 3 (maternal – orange bar) or 5 (cord – blue bar) infections per virus. Black lines indicate the sensitivity of the analysis for each virus – i.e. the efficiency that would occur if only one well across all of the infections were positive.

In order to quantify this effect, we conducted a dilution cloning experiment comparing transformation of blood from umbilical cord with blood taken at the same time from the baby’s mother. This was performed for three donor pairs, and on each occasion, both LPKO^w^ and YKO viruses consistently failed to transform cord blood, despite successfully transforming the maternal cells into LCLs (Fig 4B). In contrast, both the wild type viruses (WT^w^ and Yrev/EBV-BAC) and LPrev^i^ showed no difference in transformation efficiency between cord and maternal lymphocytes. We also observed that WT^w^-infected cells expanded and acidified the media faster than WT-HB9 and Yrev transformations, but produced the same number of LCL-initiating events (not shown). This demonstrated that the defect in transformation of cord blood is due to the EBNA-LP mutation, and is not a consequence of a generically reduced transformation competence.

In order to assess whether the cord cell phenotype was linked to the naïve phenotype of these cells, we used CD27 and IgD status to sort CD19+ve adult cells into naïve and memory B cell populations and infected them with the EBV strains. Unlike the comparison of whole adult and cord blood, we observed that for any donor, adult naïve (CD27-IgD+) B cells transformed less efficiently than either CD27+ve subset, although there was considerable variation between donors. Nine attempts were made to generate LPKO LCLs from the naïve subset of six donors, with memory cells and WT^w^ transformations as controls. Eight attempts failed to generate naïve LPKO^w^ LCLs. One donor exhibited an extremely high level of transformation by all viruses. In this case, widespread cell death was observed in the LPKO^w^-infected naïve cells 2 weeks post infection, but an LPKO LCL was established. It has previously been shown that IgD status of LCLs matches that of the initially infected cell population [37]. We therefore analyzed these LCLs for IgD and CD27 status to compare with the status of the cells as originally infected. The LPKO LCL that grew from the naïve population was clearly CD27 positive, whereas all other LCLs tested matched their original phenotype (not shown), suggesting that this LCL either arose from a mis-sorted memory cell, or somehow changed its differentiation state after sorting, but was not a naïve LPKO LCL. Overall this shows EBNA-LP is essential for the transformation of B cells with a naïve phenotype, both of cord and adult origin.

### EBNA-LP facilitates the transcription of viral but not host genes

It has been widely reported that cotransfection of EBNA-LP is able to enhance the transcription of viral and host genes induced by EBNA2 [7,22-26]. We therefore undertook qPCR analysis of host and viral transcripts for two independent time courses studying RNA levels across the first 30 days after infection of CD19 isolated B cells at an MOI of 2. The EBNA-LP mutant viruses (LPKO^i^, LPKO^w^ and YKO) all showed the similar gene regulation while the cells survived (not shown). The WT-HB9, WT^w^, E2rev, Yrev and LPrev^i^ also generally behaved the same, although LPrev^i^ sometimes diverged on day 30. The data for a representative time course are therefore presented as the comparison between these two groups (Fig 5 and Fig S10). EBNA2 transcription was assessed by a qPCR assay extending from exon Y2 to downstream of the exon Y3 splice donor. It therefore detected transcription even in the E2KO virus, which retains these sequences. EBNA2 transcript levels were very similar across all infections (Fig 5A), except that the E2KO virus had a 10-fold higher level transcript in both the EBNA2 and Wp assays (not shown), which is consistent with the elevated levels of EBNA-LP protein in E2KO infections (Fig 3C). To our surprise, however, we observed that all other virus genes tested expressed lower transcript levels in the EBNA-LP mutant group early after infection. Across a range of viral genes, expression recovers slowly to return to wild-type levels (Fig 5 and Fig S10).

**Fig 5.**
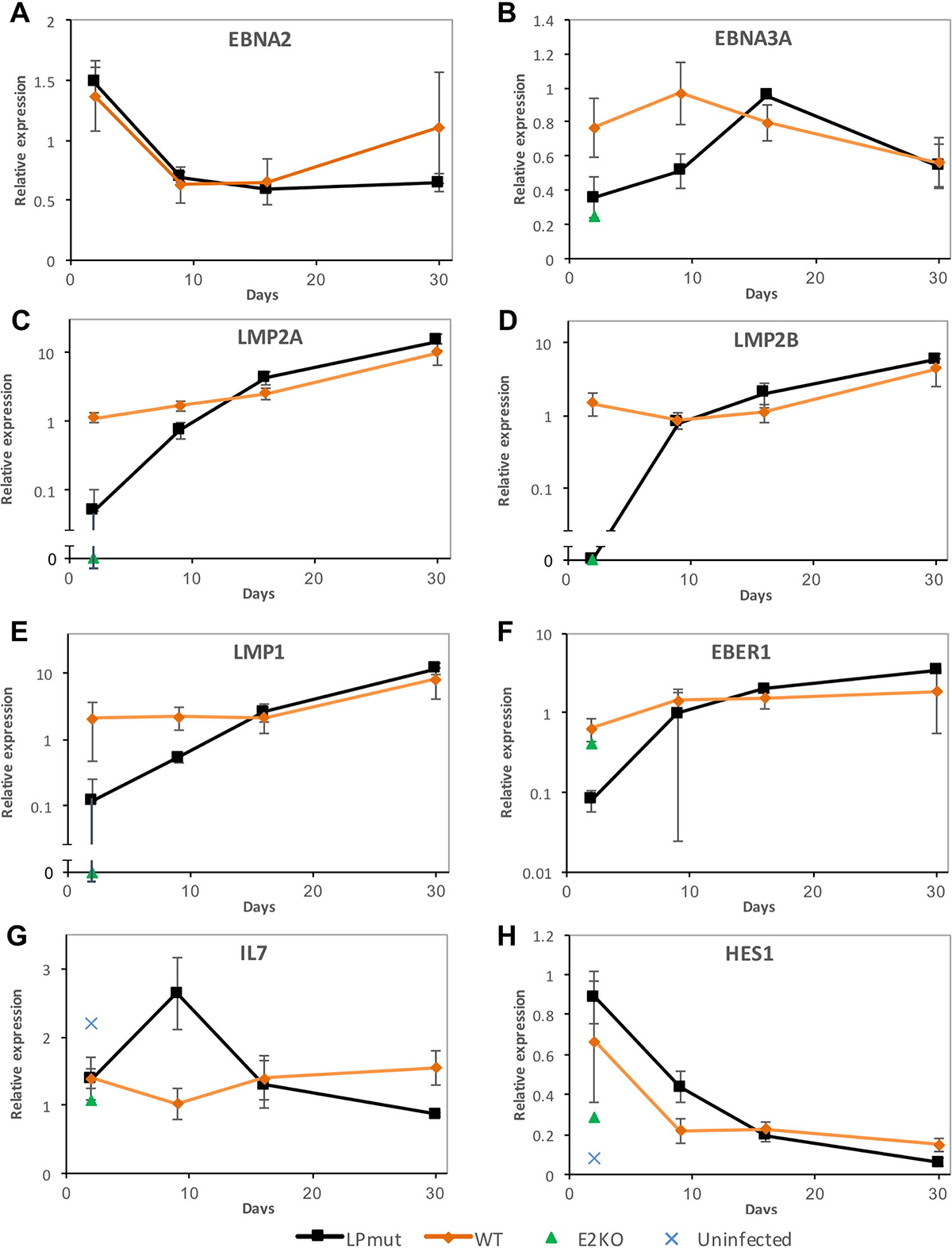
Time course for virus and host gene expression after EBNA-LP and EBNA2 mutant virus infections. Graphs show levels of virus and host transcripts as a time course after infection of resting B cells. Infections are grouped as either ‘wild-type’ (comprising WT-HB9, WT^w^, Yrev, E2rev and LPrev^i^) or EBNA-LP mutants (LPKO^i^, LPKO^w^ and YKO), since these mutants showed consistent phenotypes. These are compared with E2KO-infected and uninfected cells on day 2, as indicated by the key. Transcript levels (measured by qPCR) are expressed relative to the level for the WT-HB9 wild-type infection on day 2. Error bars show ±1 standard deviation of the gene level for each group. Note that EBNA2 transcript level in E2KO infection was >10, so is omitted. EBNA3A transcript level (B) is therefore shown relative to the EBNA2 transcript level, since they share promoters. Broken axes (C-E) are used to allow zero values to be visualised on an otherwise logarithmic axis. For virus transcripts, uninfected B cells did not show significant levels of viral transcripts (i.e. are effectively zero) so are not shown.

For EBNA transcripts, EBNA-LP makes no difference to Wp activity (Fig S10A), whereas Cp activity is generally lower in EBNA-LP mutants (Fig S10B). Transcripts for all three EBNA3s (measured across the U exon/EBNA3 splice junctions) were around one third lower early after infection with the EBNA-LP mutants. This is also true of E2KO infection, once the higher basal transcription of the Cp/Wp transcripts is accounted for (Fig 5B and Fig S10C-D).

Other viral genes were more dramatically altered in EBNA-LP mutants early after infection. EBNA2-dependent LMP2A and LMP2B-initiated transcripts, and LMP1 - Fig 5C-E) were at less than 10% of the levels seen in wild-type infections. Total LMP2 levels were generally (but not universally) lower in EBNA-LP mutants, but high in E2KO (Fig S10E), suggesting LMP2 transcription from the TR promoter may not be suppressed in these mutants. More surprising was the observation that levels of both EBV-expressed small RNAs (EBERs) – which are not EBNA2-regulated – were also much lower in the EBNA-LP mutant infections than wild-type (Fig 5F and Fig S10F). As the transformed cells grew out, the levels of all of the viral genes returned to equivalent levels between the groups, although established LPrev^i^-infected cell lines exhibited elevated levels of LMP1 and Wp transcription (not shown).

In contrast to the widespread differences in virus gene expression, EBNA2-associated host genes showed very little difference between EBNA-LP mutant and wild-type infections after 2 days. Cyclin D2, which was reportedly enhanced by EBNA-LP [7] exhibited lower transcript levels on day 9, but not consistently lower on day 2. This is likely a consequence of slower proliferation, as LPrev^i^-infected cells (which proliferate more slowly than other wild-type infections) have lower Cyclin D2 levels than other wild-types. In contrast, MYC levels were not affected by EBNA-LP (Fig S10G-H). EBNA2-dependent activation of HES1 but not CD21 was sometimes reported to be enhanced by EBNA-LP [38] [26]. IL7 has been shown to be bound by EBNA2 [14], but is not activated by EBNA2 during infection (Fig 5G). Despite these differences in reported associations with EBNA2 and EBNA-LP, all three genes show the same pattern of regulation: they exhibit a consistent increase in transcript levels only on day 9 after infection with EBNA-LP mutants (Fig 5G, Fig 5H, Fig S10I). This is the opposite effect to what might be expected if EBNA-LP contributed the enhancement of activation by EBNA2, casting doubt on this current perception of EBNA-LP function.

### EBNA-LP facilitates transcription factor recruitment to the LMP promoter

It has been reported that EBNA-LP can be detected by chromatin immunoprecipitation at various genomic loci, often in the presence of EBNA2 [39]. We have attempted to perform EBNA-LP chromatin immunoprecipitation (ChIP), but have been unable to detect any difference in EBNA-LP ChIP-qPCR signal at either LMP or Cp promoters between wild-type and EBNA-LP knockout viruses in LCLs or during primary infections (not shown). Since EBNA2 has been repeatedly shown to regulate and bind at these genes, the binding of EBNA2 to both known binding sites and negative control sites was assessed across three 30 day infection time courses. Differences in EBNA2 binding were sometimes detectable on day 2 (not shown) but the ChIP showed a much better signal to noise ratio on day 5 post infection. There is a profound delay in the recruitment of EBNA2 to known transcription factor binding sites at the LMP2A and LMP1/2B promoters on the LPKO^w^ genome compared to WT^w^ (Fig 6 and Fig S11). EBNA2 recruitment to its binding site at Cp was modestly reduced in LPKO^w^ infections, but still showed a considerable binding signal. In contrast, EBNA2 was efficiently recruited to host genes IL7 and HES1. Indeed, the LPKO^w^ infection consistently showed elevated binding on days 5 and 9, but not at other time points (Fig 6B).

**Fig 6.**
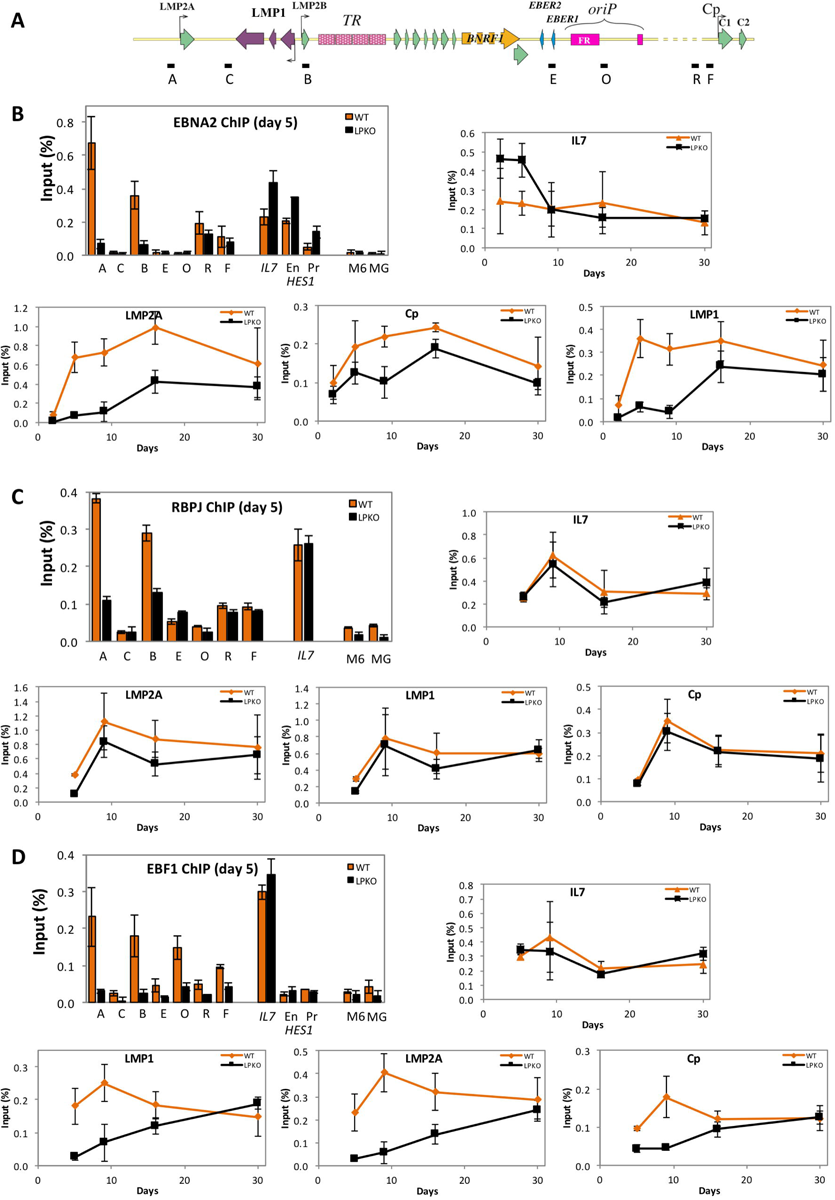
Binding of EBNA2, RBPJ and EBF1 to viral and host loci. ChIP analyses of EBNA2, RBPJK and EBF1 at promoters regulated by EBNA2 in LPKO^w^- and WT^w^-infected cells. EBV ChIP assays are shown positionally as letters in the schematic **A.** Data for ChIP of EBNA2 (**B**), RBPJK (**C**), and EBF1 (**D**) are shown for all assays on day 5 post infection with infection with WT^w^ (orange) and LPKO^w^ (black), and then for IL7, LMP1 (assay B), LMP2A (assay A) and Cp (assay R).

EBNA2 does not bind directly to DNA, but rather it is directed to many of its binding sites by host proteins, in particular the transcription factor RBPJ (also called RBP-J_κ_ and CBF1). We therefore also assessed binding to these locations. In our time course, RBPJ binding peaked later than EBNA2 binding, and was slightly (but consistently) lower in LPKO^w^ at day 5 post infection at the LMP promoters but identical at Cp (Fig 6C). However, no differences were observed at later time points, or on host genes, suggesting that the apparent lag in RBPJ recruitment to the LPKO^w^ genome is very slight.

Recent genome-wide analyses have shown that EBNA2 and RBPJ are often located with early B cell factor (EBF1) on the genome [39], and that the three proteins can bind together to chromatin [14]. In addition, EBF1 has two RBPJ-independent binding sites on the EBV genome, one near to the EBERs and the other near oriP. In wild-type infections EBF1 binding reached maximal levels at all viral sites between 5 and 9 days post infection, similar to EBNA2, whereas recruitment to both LMP and EBER/oriP loci were delayed in LPKO^w^ infection. Just like EBNA2, EBF1 recruitment to the LPKO^w^ genome was delayed, taking at least two weeks to approach wild-type levels at all of the viral locations tested. In contrast, EBF1 levels on the IL7 promoter were similar between LPKO^w^ and WT^w^ throughout infection.

Taken together, these observations showed a widespread failure of the LPKO^w^ virus to support the recruitment of transcription factors to the region of the viral genome between LMP2A and oriP, with only a slight delay at Cp. In contrast, host genes exhibited accelerated EBNA2 recruitment. These observations match the transcript level data, with the genes whose activation was most delayed also having a delayed recruitment of transcription factors. Together this suggests that EBNA-LP is required to facilitate the recruitment of transcriptional activators to the region of the EBV genome between LMP2A and oriP, but not for the activation of host genes by EBNA2.

## DISCUSSION

### We have developed a novel strategy for markerless modification of repeat regions

The genetic analysis of EBNA-LP function represents a major technical challenge, due to the repetitive and diverse nature of its gene. Transcription can initiate at either Cp upstream of IR1, or at Wp within any one of the IR1 repeat units. Previous approaches to genetically assessing EBNA-LP function have been restricted to either truncation of the protein by removal of the C terminal Y exons [27,28] or deletion of increasing numbers of W repeats, which affects Wp numbers and the sisRNAs as well as EBNA-LP [29]. Our analysis aimed to compare the function of the previously assessed C-terminally truncated EBNA-LP with a more comprehensive knockout of the EBNA-LP reading frame.

We have described two approaches to generating mutated repeat regions. The first used type IIS restriction endonucleases, based on a method for generating tandem repeats [40] that has been used for analyzing gammaherpesvirus terminal repeats to separate persistence and packaging functions [41,42], and has since been rebranded as Golden Gate cloning. This method is likely to be effective for some repeats (particularly those with many repeat units, as it can expand repeat units exponentially), but the need to mutate the BsmBI restriction site in IR1 (and the fact that this mutation appears to have detrimentally affected transformation) means that this approach is not currently useful for modifying IR1.

The second approach is novel, using Gibson Assembly [36] to seamlessly generate an array of 6 repeat units. This strategy is sufficiently controlled that it has the potential to generate any combination of IR1 repeat units in any order: mutations can be selectively generated in any of the repeat units of IR1, and the array assembled accordingly. Not only could this be used to mutate an IR1 feature in all of the repeats, as we have shown for EBNA-LP mutation, but it can also be used to specifically assess whether features play different roles depending on which repeat unit they are in (i.e. whether the roles of the first, last or internal repeats may be different). This may be important to understanding the biology of IR1, as it was reported that the first Wp in the genome may be more important than the others [43], so understanding the structure and function of IR1 will require this sort of approach.

### An intronic mutation causes a transformation defect that is independent of EBNA-LP

Our initial attempt to assess the function of EBNA-LP proved flawed: LPKO^i^ was severely defective in transformation, and the revertants (LPrev^i^) were also somewhat defective. The intronic mutations reduced the proportion of cells that divided after infection, and caused slower outgrowth. Importantly, the intronic defect was additive to the LPKO defect, indicating that the intronic mutations exerted their effect independently of transcription or EBNA-LP function.

In our view, it is most likely that the mutation deliberately introduced into the small intron (from which sisRNA1 is produced) is responsible. The other three changes, are clustered in the BWRF1 ORF, and normally exist in only one repeat unit of the B95-8 IR1 (Ba abdullah et al; manuscript submitted for publication). Two of the changes are also seen as SNPs between virus strains, so are unlikely to have a negative impact on virus fitness. All three variants produced missense changes in the putative BWRF1 ORF, although It remains unclear whether this ORF has a function, having no ATG initiation codon, no promoter, and being disrupted in around 20% of strains (Ba abdullah et al; manuscript submitted for publication). While this region is reported to persist as a stable RNA (sisRNA2), this has only been detected in the form of elevated levels of reads in RNA-seq [30]. However, sisRNA2 potentially has features in common with the alphaherpesvirus latency-associated transcript (LAT), a spliced, intron-derived RNA important for establishing latency in neurons (reviewed in [44]).

SisRNA1 is better characterized, being detectable by Northern blot, qPCR and RNA-seq [30]. Generally, sisRNAs appear to be a relatively abundant form of non-coding RNA, originally identified in Xenopus [45], but since found to be widespread in human and drosophila cells [46,47]. They have been variously proposed to regulate transcription, translation and sequester other cellular components (reviewed by [48]). Short exons closely resembling sisRNA1 have been found associated with Argonaute, and these may repress mRNA translation in a sequence-specific manner [49], which fits with the observations that sisRNA1 is detectable in oligo-dT-purified RNA [50].

It is possible that the intronic mutations could alter splicing profiles. Neither EBNA-LP nor EBNA2 transcript or protein levels were detectably altered by the intronic mutations early after infection. However some other viral gene expression may be altered. For instance, BHRF1 transcripts were recently reported to be spliced between W1 exons (skipping exon W2) during the lytic cycle, so it remains possible that this transcript, or some other at yet uncharacterized IR1 splicing pattern during early infection could be affected. But regardless of their mechanism of action, our observations represent the first evidence that the sisRNAs (or intronic sequences in IR1) may be functionally important for EBV transformation. Further study is required to establish which mutation is important, and what its functional consequences and mechanism may be.

### EBNA-LP is more important in transformation of naïve B cells

It is not unheard of for EBV mutants to exhibit different transformation phenotypes in different B cell subsets, as BZLF1-knockout EBV is better able to transform germinal center B cells than memory or naïve B cells [51]. Nevertheless, the difference between naïve and memory cells is still surprising, as (at the transcriptome level) they are much more similar to each other than to germinal center cells [52]. Since the death of LPKO-infected naïve cells was consistent for infection of both mixed lymphocyte and CD19-isolated B cells, the difference must be intrinsic to the B cell subsets.

It is enticing to invoke the differences in transformation between cord, memory and adult naïve B cells with different viruses as a possible mediator of the different characters of primary infection in individuals of different ages. The incidence of infectious mononucleosis during and after adolescence could be a result of different balances of memory and naïve cells, or differences in the character of the B cells at different ages. Since only 25% of students that seroconverted at university exhibited symptoms of infectious mononucleosis [53], the relative numbers of naïve and memory B cell, or some measure of tonsillar maturity, could be influencing the different severities of primary infection in these individuals, perhaps on a background of EBNA-LP diversity.

Biologically, a number of differences have been reported that separate the behaviour of memory and naïve cells. We noted a slower outgrowth of LCLs from naïve than from memory B cells, although this was not seen in a previous study [37]. Interestingly, adult naïve B cells entered cell cycle later than memory cells after CD40L stimulation [54], and produced fewer cells from such cultures [55]. In cord cells, the defect is more profound, with CD40 agonism barely inducing any activation markers, whereas equivalent naive B cell subsets from adults did respond [56]. Additionally, IgM crosslinking in cord cells failed to induce ERK1 phosphorylation, in contrast with adult cells [56], while BCR crosslinking on adult cells induced a larger response in memory than naïve cells [57]. These observations are most intriguing, since antibody crosslinking and CD40 activation are mimicked by the LMP proteins [58] whose expression are delayed in LPKO infections, suggesting that perhaps the deregulation of LMPs in LPKO may be responsible for this effect, so it would be intriguing to investigate how LMP1 and LMP2 knockout EBVs behave during transformation of naïve B cells.

Other phenotypic differences between naïve and memory cells may also contribute. For instance, IL2 stimulation enhanced the production of memory cells by CD40L, but not naïve cells [54], while 95% of cord B cells are negative for the IL2 receptor [56,59]. Naïve B cells also have a much lower level of the anti-apoptotic bcl2 family members MCL-1 and Bcl-x_L_ (but not Bcl2 or Bim) than memory cells [60,61], which may contribute to the apoptotic phenotype of the LPKO^w^ naïve cells.

Together these reports suggest that naïve and memory B cells are phenotypically different, both in their response to pro-proliferative signaling and their resistance to apoptosis. What is less clear is how EBNA-LP overcomes these differences in naïve cells. It has been reported to interact with a complex of tumor suppressors MDM2, p53 and the cyclin-dependent kinase inhibitor p14^ARF^ [62], which could influence both proliferation and apoptosis responses. Alternatively, metabolic stress has been reported to be an important limitation to B cell transformation, and appears linked to an elevated EBNA-LP:EBNA3 ratio [63]. Furthermore, both EBNA-LP and EBNA3A have been shown to bind the prolyl-hydroxylase proteins that influence HIF1a stability, with the suggestions that this alters the metabolic state of the infected cells [64]. However, further study is required to understand the biology underlying the difference in transformation of naïve and memory cells, and to understand whether these differences are important for the in vivo biology and pathogenesis of EBV.

### EBNA-LP only enhances the transactivation of genes by EBNA2 in specific circumstances

The function of EBNA-LP has been linked to EBNA2 because of their co-expression immediately after infection, and from a series of co-transfection experiments that appeared to show an enhancement of EBNA2’s transactivation function in the presence of EBNA-LP [7,22,23,25,26,38,65]. These studies demonstrated an ability to enhance transcription from reporter constructs [23,65], from host genes - most notably *HES1* and *CCND2* (Cyclin D2) [7,38] - and from EBV promoters repressed in the latency I transcriptional profile, including *LMP1* [22,26], *Cp* [65] and *LMP2A* [38]. Some studies have failed to replicate the regulation of some of these genes [26], but activation of the LMP1/LMP2B bidirectional promoter has been observed consistently. Our approach differs considerably from the previous ones, both by taking a genetic approach to controlling the presence of EBNA-LP, and by analyzing gene expression in the context of viral infection rather than using isolated EBV proteins. Overall we have observed a much more widespread than expected impact of EBNA-LP on viral gene expression after infection, which contrasts with a delayed and transient impact on host gene transcript levels.

Of the previously studied host genes, we have seen no conclusive effect of EBNA-LP on *CCND2* transcription, in contrast with a previous report [7]. Notably the differences in cellular proliferation between 5 and 14 days post infection is more likely a cause than a consequence of the transcript differences in CCND2 seen on day 9 (Fig S10), as it is also seen in LPrev^i^. For other EBNA2-induced host genes we have seen transient but highly reproducible increases in both transcript levels and binding of transcription factors (EBNA2 and EBF1) to the genes at day 9 post infection with LPKO viruses. Notably this is the opposite to what is predicted by the EBNA2-enhancement hypothesis espoused by the previous literature. This increased transcription and EBNA2 binding in LPKO^w^ could reflect a genuine co-regulation of the host gene by EBNA-LP and EBNA2. ChIP-seq analysis of EBNA-LP binding to the genome has suggested that it can be found associated with EBNA2 [39], and EBNA-LP has been reported to bind to EBNA2, albeit only when its acidic C-terminus is deleted [66]. Nevertheless, the transient increase in EBNA2 binding in LPKO^w^ infection could simply be a consequence of excess availability of the EBNA2 that failed to bind to the viral genome. Either way, these data clearly show that EBNA-LP does not enhance the transactivation of host genes by EBNA2, as previously claimed, but can contribute to EBNA2 recruitment to host genes.

### EBNA-LP has a widespread effect on EBV gene expression

In contrast to host genes, widespread viral transcription is profoundly delayed in the absence of EBNA-LP. The EBNA transcripts are all generated from alternative splicing after transcription initiation at either Wp or Cp promoters. While EBNA2 levels were not affected by the loss of EBNA-LP, suggesting no change in promoter activity, the reduced levels of the EBNA3 transcripts downstream suggest that the processing of the transcripts is different in the LPKO infection. This elevated ratio of upstream to downstream EBNAs (as a ratio of EBNA-LP to EBNA3C protein levels) was observed in cells that have only proliferated 1-3 times after EBV infection [35], and in cells that arrest after an initial period of proliferation [63]. This could result from either an increase in polyadenylation after EBNA2, a change in splice site usage, or reduced elongation of transcripts. Indeed, there is evidence that the elongation complex pTEFb is important for transcriptional elongation from Cp, but is predicted to be less important for Wp [67], leading to speculation that elongation of Wp transcripts is less efficient, which would lead to lower yields of downstream EBNAs. We have seen reduced levels of downstream transcripts, along with modestly delayed EBNA2 recruitment to and transcription from Cp in EBNA-LP mutant viruses, which could explain this phenomenon.

A more profound effect was seen on the EBV latency genes between LMP2A and oriP (see schematic in Fig 6A). Activation of this whole genome region was severely delayed in LPKO infections, and this correlated with the delayed recruitment of EBF1, EBNA2 and - albeit less dramatically - RBPJK. The failure to induce transcription of the EBERs demonstrates that EBNA-LP is not simply working through EBNA2, and raises the question of whether using EBER in situ hybridization to diagnose EBV-positive malignancies is reliable in all contexts.

The region of latency genes from the LMP2A promoter to oriP represents a coordinately regulated genomic locus. It is flanked by CTCF binding sites [68], and these loop together to form a transcriptional unit. Disruption of the CTCF site near the LMP2A promoter can disrupt this loop, consequently reducing LMP gene transcription and increasing repressive histone and DNA methylation in LCLs [69]. The simplest interpretation of our data is that EBNA-LP is important for the proper establishment of this transcriptional unit. By 4 weeks post infection, the LPKO LCLs have reached normal expression levels of LMPs and EBERs, so there does not appear to be a defect in the maintenance of the locus once it is established. However, there is a profound delay in the recruitment of transcription factors. Indeed, EBF1 and RBPJ have been described as a pioneer factors: transcription factors that are able to access chromatinized DNA and establish new enhancer regions [14]. However, while EBNA2 and EBF1 are readily able to access cellular loci in the absence of EBNA-LP, they appear to require it to efficiently access the incoming EBV genome.

### Possible mechanisms of action of EBNA-LP

The ability of EBNA-LP to enhance EBNA2-dependent gene transcription has been variously attributed to its ability to bind to Sp100, HDACs 4 and 5 [65], or NCOR [38]. The binding to Sp100 is interpreted to transiently disrupt PML nuclear bodies (ND10) early during infection, and thereby evade an as yet undefined antiviral process [21]. EBNA-LP binding to NCOR and HDACs are both reported to sequester these repressive proteins away from EBNA2-inducible genes, thereby improving transactivation [38,65]. Any (or a combination) of these remain reasonable hypotheses as to how EBNA-LP facilitates viral transcription after infection.

The major insight that we offer is that the role of EBNA-LP is tied to the transcription of the incoming DNA. This could involve evasion of the antiviral effects of ND10, which may be responsible for suppressing transcription of the incoming genomes. Additionally, retroviral genomes that fail to integrate into the host genome exhibit increased gene expression in the presence of HDAC inhibitors [70,71], supporting the idea that inhibition of HDACs by EBNA-LP could also relieve repression of the incoming viral genome. If such repression were mediated by HDACs (and perhaps also NCOR), this suggests that EBNA-LP disrupts this process at several levels.

The chromatinization of viral genomes is usually very rapid, and an EBV tegument protein - BNRF1 - has been identified that binds to Daxx (an ND10 component) and supports histone loading onto incoming genomes [72]. However, an elegant study has shown that BNRF1 and EBNA-LP have complementary effects in preventing the suppression of herpesviruses, having a combinatorial effect in helping an ICP0-null herpes simplex virus to evade the effects of ND10 [73]. It is tempting to speculate that transcription of Wp is supported by the action of BNRF1, and the EBNA-LP produced from those transcripts then prevents innate processes from inhibiting the LMP/EBER/oriP/Cp region of the genome. Considerable experimental effort will be required to test these hypotheses.

Of course, there are other aspects to this genome region that could explain why its regulation is not like that of Wp/Cp. Notably, this region includes the terminal repeats, and the virus – linear in the virion – needs to recircularize before LMP2 can be transcribed, and perhaps before this region is properly regulated. In addition the terminal repeats contain a binding site for PAX5, which is directed to the viral genome by EBER2 [74]. Two of the factors reported to bind to the EBER2/PAX5 complex (NONO and SPFQ) have also been reported to bind to EBNA-LP in a tandem affinity mass spectrometry experiment [75], although these two proteins both have a high background signal in such experiments according to the CRAPome repository [76]. Nevertheless, it is possible that EBNA-LP is involved in PAX5 recruitment to the terminal repeats.

The observation that the truncated EBNA-LP in the YKO cells localizes to the nucleolus suggests a role of this compartment in EBNA-LP function. Certain stimuli have been reported to induce nucleolar relocalization of EBNA-LP, probably through interaction with HSP70, or p53 complexes [77,78]. However, the relevance of this remains obscure. Interpreting the phenotype of the YKO viruses is difficult, as this mutant also contains the previously unreported STOP codon in one W1 exon. This may have contributed to the very low level of truncated EBNA-LP, and this level may be low enough for the virus to be functionally null for EBNA-LP in some of the biological readouts. Previous analyses of EBNA-LP truncated viruses either did not report analysis of EBNA-LP protein levels [27], or failed to detect it [28], although it is unclear whether their antisera could detect the type 2 EBNA-LP of the parental P3HR1 virus. Nevertheless, YKO was less defective in the initial transformation and proliferation, despite showing the same apoptotic phenotype in naïve cells, suggesting that the Y domains are crucial for this latter biological function.

In summary, we have undertaken a genetic analysis of EBNA-LP function and shown that EBNA-LP is important for B cell transformation, and essential for the transformation of naïve B cells, and that the role of EBNA-LP is far more complex than the previously proposed cofactor for EBNA2, being particularly important for establishing the viral transcription program. We also suggest that future analyses of EBV mutants would be better performed in distinct B cell subsets, as it is clear that phenotypes can vary considerably according the differentiation state of the infected B cells, and perhaps also the age of the B cell donor. The observations and genetic manipulation strategies described herein also extend approaches to study EBNA-LP, the EBV-sisRNAs and the wider functions of IR1 in the future.

## METHODS

### Generation of recombinant EBVs

In order to introduce mutations into IR1, we have devised a strategy for introducing a constructed IR1 repeat into EBV. This entails first deleting the virus’s endogenous IR1 (to prevent the constructed repeat from recombining with the original one) and then inserting the rebuilt repeat. To achieve this, we used RecA-mediated recombineering as previously described [79]. The viral IR1 was deleted by joining together homology regions from the unique (non-repetitive) sequences flanking IR1: The upstream region (NC_007605 positions 11413-12008), which contains exon C2, was cloned SfiI/PciI from the B95-8 BAC (clone WT-HB9); the downstream region (position 35239-35869) was cloned XhoI/MluI. This region was introduced by recombineering in place of IR1. The same homology regions were used as flanks for newly assembled IR1 repeats containing EBNA-LP mutations.

We have used two distinct methods to generate the synthetic IR1. Both approaches generate an IR1 with 6.6 copies, which is a typical size for circulating EBV strains [31] and is the size of IR1 in the parental EBV-BAC clone, WT-HB9. In both cases, the IR1 was assembled in a pBR322-based plasmid in DH5alpha bacteria grown at 30°C to reduce unwanted recombination.

The first approach used to assemble a modified IR3 adapted a strategy that used type IIs restriction endonucleases to assemble repeats [41]. A BamW fragment was subcloned from the B95-8-BAC clone WT-HB9 into a vector that contains binding sites for the type IIb restriction endonucleases BsmBI and BtgZI. These restriction sites were engineered to both cut at the site of the BamHI restriction site (Fig S1A). A DNA fragment (between the MfeI and AgeI restriction sites in BamW) was synthesized, containing a point mutation of the BsmBI restriction site in the intron between exons W1 and W2, and also containing mutations that introduced STOP codons and a PvuI restriction site, for making the EBNA-LP knockout virus, LPKO^i^. A second synthesized fragment containing the BsmBI mutation but not the EBNA-LP mutation was also synthesized for producing the revertant virus, LPrev^i^. These fragments were cloned into the BamW repeat unit, and then both the LPKO^i^ and LPrev^i^ repeat units were assembled into an array using the method described in Fig S1B. The array was then incorporated into independent IR1 knockouts [WKO] according to the scheme shown in Fig S1C, generating two independent LPKO^i^ viruses, and their revertants.

Subsequently, recombinant viruses were made that contained changes in IR1 without need to mutate the BsmBI restriction site in the W1-W2 intron. This was achieved by cloning a new BamW repeat unit that matched the B95-8 consensus sequence into a pBR322-based plasmid that contained BtgZI restriction sites that cut the BamHI sites flanking the repeat unit. This was then modified with oligonucleotide linkers on either (or both) sides of the BamW fragment, such that the BamW sequence was extended approximately 20bp from the BamHI restriction site (Fig S8A). Additional constructs were generated by cloning each of the flanking regions (described above) adjacent to the BamW fragment. To generate the wild-type IR1, the constructs were cut and assembled as shown in Fig S8A, and the assembly was cloned into pKovKan and recombined into the WKO.4 that had been used to produce LPKO^i^.2, thereby generating WT^w^.1 (see Fig S1C). To generate the new EBNA-LP knockout (LPKO^w^) the BsmBI point mutation in the synthesized LPKO region was reverted to the wild-type sequence by InFusion mutagenesis, and subcloned into the new wild-type BamW fragment. The IR1 synthetic array was then assembled in the same way as the wild-type array, and used to independently generate LPKO^w^.2 and LPKO^w^.4 viruses by recombineering into WKO.4. E2KO, E2rev, YKO and Yrev BACs were generated by RecA-mediated recombineering essentially as described elsewhere. The precise sequences of the E2KO and YKO deletions are shown in Fig 1. Revertants were made by reintroducing wild-type sequence into the knockouts by the same method (Fig S1).

BACs were screened for integrity using EcoRI, AgeI, HindIII, NotI and BamHI restrictions digests and run on a CHEF DRII chiller pulsed filed gel electrophoresis system (Bio-Rad). We noted that the family of repeats (FR) region of oriP is smaller in WT-HB9 than predicted by sequence. This reflects a previous observation that the family of repeats region (FR) of oriP is unstable, even in BACs, and that the FR in the p2089 BAC (of which WT-HB9 is a subclone) is 300 bp smaller than the authentic sequence of B95-8 [80]. Therefore, in addition to restriction digests, all recombinant BACs were screened by PCR, using the KA2 and KA3 primers [81] with Q5 DNA polymerase (NEB) to ensure that the FR region was the same size in all recombinants.

### Generation of producer cell lines and virus

Recombinant EBV BAC DNA was purified from bacteria by alkaline lysis followed by cesium chloride density gradient centrifugation. DNA was assessed by pulsed field gel electrophoresis to ensure a predominance of intact supercoiled BAC DNA, as DNA integrity appears to influence the number and quality of producer cell lines. The BAC DNA was transfected into 293-SL cells (a culture of the HEK-293 cell line provided by Claire Shannon-Lowe; University of Birmingham) using a peptide 6 and lipofectin transfection reagent described previously [82]. Cells were selected with hygromycin and colonies isolated by ring cloning. Individual hygromycin-resistant colonies were screened for GFP expression, for their ability to produce virus. The integrity of episomes from the producer lines was assessed by recovery into bacteria [83] and analyzed by restriction digest and pulsed field gel electrophoresis. Cell lines were used if at least 80% of recovered episomes were indistinguishable from the parental BAC.

To generate virus stocks, 293-EBV producer cell lines were seeded in 10 cm dishes and after 1-2 days these were transfected at approximately 25% confluency with equal quantities of BALF4 and BZLF1-experessing plasmids - 12 μg total DNA per 10 cm plate when transfecting with peptide6+lipofectin or 6 μg per plate using GeneJuice reagent (Merck-Millipore). Supernatant was harvested after 5 days and filtered through a 400 nm syringe filter. Virus titer was assayed by infecting 2x10^5^ Raji cells in 1.5 ml with 10-fold dilutions of virus. After two days, the Raji cells were treated overnight with 20 ng/ml TPA and 5 mM sodium butyrate to enhance GFP expression in the infected cells. Cell clumps were dispersed by pipetting and total number of green cells per well were counted under a fluorescence microscope. This gave a Raji green units (rgu) titer, which was typically in the range of 0.5-10x10^5^ rgu/ml in the cell culture supernatant.

### Cell culture, isolation of immune cells and virus infections

LCLs, BL31 cells (provided by Alan Rickinson, University of Birmingham), and 293-SL cells were grown in RPMI media supplemented with L-glutamine (Life Technologies) and 10% fetal calf serum. This serum was batch tested for the ability to establish 293-SL-EBV-BAC colonies after BAC transfection, and to support outgrowth of LCLs under limiting dilution. MRC5 foreskin fibroblasts (ATCC CCL-171), also grown in RPMI, were irradiated with 50 Gy and seeded as a confluent monolayer to support outgrowth in some experiments.

Adult primary lymphocytes were isolated mainly from buffy-coat residues, but also from lymphocyte cones, both provided by NHS Blood and Transplant. Cells from a 500 ml original blood volume were diluted to 200 ml with PBS. Lymphocytes were isolated by layering blood-derivative on ficoll followed by centrifugation. The isolated peripheral blood leukocytes (PBLs) were washed twice in RPMI/1%FCS. B cells were purified from PBLs by hybridizing to anti-CD19 microbeads (Miltenyi), using 0.5ml beads per 10^9^ PBLs, followed by positive selection (possel program) on an autoMACS separator (Miltenyi). Either purified B cells or PBLs were resuspended at 1-2x10^6^ cells/ml in RPMI/15% FCS. B cell purity was measured by FACS for CD20 positivity, and was typically around 95%.

For isolation of different adult B cell subsets, the CD19-sorted B cells were rested overnight in a cell culture incubator, and then stained with fluorescent antibodies (from Biolegend) against IgD (PE-CF594, clone IA6-2) and CD27 (PE-Cy7, clone M-T271). The cells were sorted using a BD FACSAria III (BD Biosciences) into three populations: naive (IgD^+^CD27^-^), class-switched memory (IgD^-^CD27^+^) and unswitched memory (IgD^+^CD27^+^). Cell populations were counted and resuspended in RPMI/15% FCS at 2x10^6^ cells/ml.

Isolated PBLs or B cells were infected within a few hours of isolation/purification, by adding virus at an MOI of 1-2 rgu/B cell, and shaking at 37°C for 3 hours, after which cells were centrifuged at 200g for 10 minutes and seeded at a density of 1-2x10^6^ cells/ml in RPMI supplemented with L-glutamine and 15% FCS (batch tested for LCL outgrowth – GE healthcare or Life Technologies) and either 50 ng/ml (for purified B cells) or 500 ng/ml (for mixed lymphocytes) of cyclosporin A. During outgrowth, approximately half of the media volume was replaced every 5-7 days (cyclosporin A was omitted after two weeks), harvesting up to half of the cells, depending on experiment.

### Transformation assay for maternal and cord umbilical blood

Blood from the umbilical cord and maternal blood was drawn from healthy full-term pregnancies. Mononuclear cells were isolated from paired 0.5-2 ml blood samples of maternal and cord blood by ficoll gradient centrifugation. Variations in the yields of mononuclear cells meant that different infections were performed with different numbers of cells: two of the three donors used equal cell numbers for maternal and cord blood infections (3.4x10^5^ and 1x10^5^ cells per infection). The third pair used 5x10^4^ maternal cells, and triplicate infections of 3x10^5^ cord cells for each virus. For most viruses (LPKO^w^.4; WT-HB9; WT^w^; LPrev^i^; YKO.4 and Yrev.4) 10^5^ Raji infectious units were used for each dilution series. LPKO^w^.2 was used at 10^6^ rgu per dilution series, but this higher titer showed the same transformation efficiency as LPKO^w^.4.

Each infection (and an uninfected control well) was placed in a well of a 96 well plate, and then serially diluted 2-fold ten times in RPMI/15% FCS/Cyclosporine A (100ng/ml). Media was changed weekly and after 6 weeks the number (n) of wells containing LCLs was counted, and number of transforming events per infection calculated as 2^(n-1)^.

### RNA analyses and quantitative reverse transcript PCR

For the time courses after infection of primary B cells, the cells were supplemented with an equal volume of fresh media 24 hours prior to harvesting. Then, half of the culture was taken (typically 5x10^5^ to 2x10^6^ cells) and RNA was extracted using RNeasy mini columns (Qiagen). For all samples in a time course, the same quantity of RNA (~300ng) was reverse transcribed using either Superscript III First-Strand Synthesis SuperMix for qRT-PCR (Life Technologies) 3 μl cDNA was mixed with TaqMan gene expression mastermix (Life Technologies) applied to a custom TaqMan low density array (TLDA) card containing duplicate assays (table ST1), which used ALAS1, RPLP0, GNB2L1 and 18S RNA as endogenous control genes. EBV TaqMan assays were designed by Applied Biosystems/Life Technologies using proprietary software, and validated using B95-8 cDNA. Sequence information is proprietary. The assay IDs in table ST1 can be used to obtain these assays. The EBNA3 TaqMan assays were designed spanning the exon junction between the U exon and the first exon of each EBNA3. LMP exon junctions detected by LMP assays are shown in table ST1. Additional assays (primers in table ST2) were conducted using Kapa qPCR SYBR kit (low ROX), and the IL7 TaqMan assay used Takyon low ROX Probe 2X MasterMix dTTP (Eurogentec) and normalised against ALAS1 and RPLP0. Quantitation of qPCR data was performed using the delta-delta-Ct method, using DataAssist Software v3.01 (Thermo Fisher Scientific). All quantitation is expressed relative to the level for WT-HB9 on day 2 post infection. Bulk PCR of transcripts across IR1 was performed using Q5 polymerase (NEB) and Cp-forward or Wp-forward primers with U-reverse or Y2end-reverse primers (Table ST2).

### Chromatin immunoprecipitation (ChIP)

ChIP was carried out using the Chromatin Immunoprecipitation (ChIP) Assay Kit (Millipore) according to manufacturer’s instructions. Briefly, 2x10^6^ infected B cells were fixed for 10 minutes in 1% formaldehyde and neutralised with glycine. After two PBS washes, cells were lysed with SDS Lysis buffer on ice for 10 minutes and sonicated using the Diagenode UCD-200 Bioruptor for 15 minutes. Precleared chromatin, using 45μl protein A agarose beads was diluted with ChIP dilution buffer and incubated overnight with primary antibodies against EBNA 2 (Abcam ab90543), EBF1 (Millipore AB10523), RBPJk (Abcam ab25949) or an IgG control (Sigma). Protein A agarose beads collected the immune complexes, which were subsequently washed in low salt, high salt, lithium chloride and twice in TE buffers. The immune complexes were eluted from the beads using elution buffer and left overnight at 65 degrees. After proteinase K treatment for 2 hours at 50 degrees, DNA was then purified using the Qiagen QIAQuick gel extraction kit, and eluted in 120 μl water.

Chromatin was quantified by qPCR using the Kapa qPCR SYBR kit (low Rox) on a QuantStudio7 real time PCR machine (Applied Biosystems). Primers used for ChIP have been described previously [84] [85] [14] [86] [67], and are listed in table ST3. Absolute quantity (relative to input) was calculated from standard curves generated from input DNA that was serially diluted 1:4, four times. 2 μl of ChIP sample was amplified in triplicate for each qPCR assay.

### Proliferation assay (cell trace)

Prior to infection, primary cells were resuspended at 10^6^ cells/ml in PBS containing 5 μM CellTrace Violet (Life Technologies) and incubated for 20 min at 37°C in dark. This was then diluted 5 times in complete B cell media and incubated for 5 min at room temperature in the dark. Cells were washed by centrifugation and resuspended in fresh pre-warmed complete B cell media for infection. We noted that CellTrace violet staining had a variable propensity to kill primary B cells, so individual tubes were tested for toxicity by staining PBLs and comparing B cell percentage with and without staining. Tubes exhibiting less than 50% loss of B cells were used in experiments. For assay, cells (a volume equivalent to 10^6^ cells in the initial infection) were harvested on ice and stained for CD20-PEVio770 (Life Technologies), and resuspended in PBS/1%BSA containing DRAQ7 live/dead cell stain (BioStatus). Cells were analysed on a FACS machine (BD LSR II or LSRFortessa) and cell proliferation visualised for live CD20^+^ singlet cells using FlowJo software.

### DNA fragmentation assay

Approximately 10^6^ infected B cells were resuspended in 50 μl PBS and added to 450 μl of ice cold 70% ethanol and stored until all samples had been harvested (24 hours to 7 days). Cells were pelleted by centrifugation at 500g for 5 minutes, stood in 1 ml PBS for 1 minute, pelleted, and resuspended in 100 μl PBS containing 1% triton X-100 and 1μg/ml DAPI. 30 μl of cell suspension was transferred to a NC-Slide A2 and imaged in a nucleocounter NC-3000 (Chemometec)

### Western blotting and immunofluorescence

Western Blotting was performed as described previously, using RIPA lysates and run and blotted onto nitrocellulose using the mini-Protein systems (Bio-Rad). Antibody clones used were: EBNA-LP (clones JF186 or 4D3); EBNA2 (Clone PE2); EBNA3A (Ab16126, Abcam); EBNA3B (Rat monoclonal 6C9 [17]); EBNA3C (mouse monoclonal A10); LMP1 (monoclonal CS1-4, Dako). For immunofluorescence, cells were grown on a 12 chamber slide (Ibidi). Cells were gently washed with PBS and then fixed with 4% paraformaldehyde for 15 minutes. Cells were washed twice with PBS and covered with blocking buffer (PBS/10% FCS/100mM glycine/0.2% Triton X-100) for 30 minutes. Cells were stained with primary antibody in 50 μl blocking buffer for one hour, washed thrice in PBS and stained with fluorophore-conjugated secondary antibody (Cheshire Bioscience) for an hour. Chambers were washed three times with PBS and then the chamber removed, the slide briefly dipped in deionized water, and a coverslip mounted on the slide with Prolong Gold Antifade mount with DAPI (Life Technologies). Slides were imaged on a Zeiss LSM5 Pascal confocal microscope: 63x objective, 4x digital zoom and shown as a projection of z-stacks of 1μm sections.

### Ethics statement

Adult blood cells were purchased from UK National Blood and Transplant as waste products of platelet isolation. As they are waste products from anonymous volunteer donors, no ethics approval is required. Umbilical cord blood (and the maternal blood) were obtained with written informed consent of the mother (an adult) prior to the onset of labour, under the MatImms study, approved by the UK National Health Service Research Ethics Committee (approval REC 13/LP/1712). Anonymized blood samples surplus to the requirements of the MatImms study were used in this project, distributed by the Imperial College Healthcare NHS Trust Tissue Bank (REC 12/WA/0196) and approved by the tissue bank’s Tissue Management Committee (project R15029). Other investigators may also have received these same samples.

## Supporting figure legends

**Fig S1. Schematic representations of the recombinant viruses used in this project. A.** Method for the construction of LPKO^i^ and LPrev^i^ viruses. Type IIS restriction enzyme sites used to assemble repeat arrays were designed to cut at the same site as BamHI in a pBR322-based plasmid. The BamHI sub-fragment (BamW) was subcloned into the BamHI site in the orientation indicated. The internal BsmBI restriction site that was mutated to allow this construction method is outlined by a green box. Other features of the IR1 repeat are indicated. **B.** Cloning strategy for the assembly of LPKO^i^ is shown. LPrev^i^ assembly followed an equivalent series of cloning steps. Grey boxes indicate BamW fragments, while white boxes indicate the SfiI/BamHI or BamHI/MluI regions at the edges of IR1 as shown in A above. Black box within BamW represents the mutation of EBNA-LP and the deleted BsmBI restriction site. Plasmid IDs are indicated. C and Y indicate the exons at the flanks of the targeting region. **C.** Schematic representation of the set of recombineering steps used to generate the recombinant EBVs constructed for this study. Identities of viruses as used in the text are in the larger font. Below, alternative lab names are included for reference. Coloured names indicate recombinant BACs that were used to generate the viruses used in experiments – Green names are wild-type in sequence and phenotype; Red names are mutants; LPrev^i^ is shown in purple, as it contains a point change compared to wild-type that was intended to be phenotypically neutral.

**Fig S2. Pulsed field gel analysis of recombinant EBVs.** Analyses show the diagnostic digests for the construction of: **A.** LPKO^i^ and its revertant LPrev^i^; **B.** E2KO and E2rev; **C.** YKO and Yrev. The size standard marker (M) is a 1:1 mixture of BstEII-lambda and Lambda mono-cut marker (NEB). **A.** Recombinant LPKO^i^ and LPrev^i^ viruses are identical, including all containing 6.6 IR1 repeats, other than bands altered by the inserted PvuI restriction site or removal of BsmBI. Digestion at these sites results in conversion of the IR1 band (white arrow) into the 3kb IR1 repeat unit (green arrow) and the Cp and Y bands flanking the repeat (yellow arrows). **B.** Size changes in E2KO result from introduction of EcoRI and PvuI restriction sites. **C.** YKO mutation produces a 140bp reduction in band size that is too small to detect in these digests, and an introduced EcoRI restriction site that causes a more easily observed change (red arrows). All other bands are unchanged, demonstrating the integrity of the genome outside the intended mutations.

**Fig S3. Western blot validation of EBNA2 knockouts in BL31 cells.** Various western blots for EBV proteins in cell lines infected with EBNA2 knockouts and revertants. Each lane is identified by the virus recombinant, above the identifier of the 293 cell virus producer line, and bottom is the BL31 cell line ID. Each lane therefore represents an independent cell line. Note that BL31-E2KO-GK is cell line generated using a different EBNA2-knockout EBV by Gemma Kelly and Alan Rickinson [32].

**Fig S4. EBV transcript validation in BL31 cells.** To test whether the splicing of EBNA transcripts had been affected by the changes inserted into the viruses, PCRs were conducted between the C1 and W0 exons (upstream) and the YH exon downstream to compare the transcripts produced by wild-type EBV and the LPKO^i^, LPrev^i^, and YKO EBVs. Use of a U exon primer downstream, and transcript analysis in 293-SL producer cell lines gave similar results (not shown).

**Fig S5. Transformation of B cells by recombinant viruses.** Photographs of the accumulation of transformed cells after infection of CD19-purified B cells by various EBV strains, taken on days 2-10 post infection as indicated. Activated cells form clusters that then proliferate to differing extents.

**Fig S6. Induction of proliferation by recombinant viruses.** Flow cytometry plots from live CD20-positive cells harvested either **A.** 3 days or **B.** 5 days after infection of adult B cells stained with CellTrace Violet prior to infection. Degree of dilution of the violet signal is indicated on the x-axis, indicating number of cell divisions. Proliferation of infected cells was measured by dilution of CellTrace violet. Data for day 7 are found in Fig 2B.

**Fig S7. Schematic showing differences from the consensus B95-8 sequence in the W repeat unit used to generate LPKO^i^ and LPrev^i^.** The BamW repeat unit used to construct the LPKO^i^ and LPrev^I^ IR1 repeat arrays was subcloned from the B95-8 BAC, but later found to contain several point changes relative to the previously published sequence of B95-8. These changes exist in one repeat unit in B95-8 (Ba abdullah et al; submitted for publication) and this repeat unit was unintentionally used to produce LPKO^i^ and LPrev^i^. This BamW repeat is indicated by the bracket below the schematic, and is repeated 6 times in the repeat array. Non-consensus nucleotides in BamW are indicated by a base followed by the consensus base in brackets, which is green where the non-consensus base is found as a polymorphism in other virus strains. The red base (T) creates a STOP codon in the subcloned repeat unit, but was replaced by the consensus G as a result of the LPKO^i^ and LPrev^i^ cloning strategies (indicated by →**G)**. The nucleotides at the Cp and Y exon ends of the repeat is the same as the parental BAC (the identity of G/T has not been determined). The intronic point mutation (in purple) is the one deliberately introduced into the LPKO^i^ and LPrev^i^ viruses to remove the BsmBI site (Fig 1A).

**Fig S8. Schematic showing methods used to generate repeat arrays. A.** Schematic representation of the Gibson assembly strategy used to generate LPKO^w^ and WT^w^. Grey boxes represent the BamW fragment and white boxes the flanks of the repeat as described in Fig S1. Red and orange arrows indicate the sequences either side of the BamHI restriction site within IR1. These arrows are the homology regions whose overlap drives the Gibson assembly of overlapping fragments as indicated in the lower part of the figure, which shows the assembly of wild-type BamW fragments into the IR1 used to generate WT^w^. To generate LPKO^w^, the mutated W exons were cloned into each of the five plasmids indicated, and the assembly performed in the same way. **B.** Pulsed field gel analysis of the recombinant WT^w^ and LPKO^w^ viruses compared to the parental EBV-BAC (WT-HB9). The PvuI digest shows the presence of the knockout-specific mutation in EBNA-LP (yellow arrows), releasing multiple copies of the 3kb IR1 repeat unit (white arrow), as compared to the parental BAC (WT-HB9) and WT^w^. The other digests show the overall integrity of the rest of the virus genome.

**Fig S9. Proliferation of cell lines at various time points.** Flow cytometry plots from live CD20-positive cells harvested either 4, 11 or 15 days after infection of adult B cells stained with CellTrace violet prior to infection. Degree of dilution of the violet signal is indicated on the x-axis, indicating number of cell divisions. Proliferation of infected cells was measured by dilution of CellTrace violet. Data for day 8 are found in Fig 3A.

**Fig S10. Time course for virus and host gene expression after EBNA-LP and EBNA2 mutant virus infections.** As for Fig 5, graphs show levels of virus and host transcripts as a time course after infection of resting B cells. Infections are grouped as either ‘wild-type’ (comprising WT-HB9, WT^w^, Yrev, E2rev and LPrev^i^) or EBNA-LP mutants (LPKO^i^, LPKO^w^ and YKO), since these mutants showed consistent phenotypes. These are compared with E2KO-infected and uninfected cells on day 2, as indicated by the key. Transcript levels (measured by qPCR) are expressed relative to the level for the WT-HB9 wild-type infection on day 2. Error bars show ±1 standard deviation of the gene level for each group. Note that Wp transcript level in E2KO infection was >10, so is left off the graph to allow the other data to be more clearly visualized. EBNA3B and EBNA3C transcript levels (C, D) are therefore shown relative to the EBNA2 transcript level, since they share promoters. Note that error bar for LMP2 levels (E) in wild-type infections on day 2 extends beyond 0, so is not plotted.

**Fig S11. Binding of EBF1 to viral loci that do not bind EBNA2.** Time course of ChIP analyses of EBF binding to its sites near the EBERs and oriP. These are respectively assay E and assay O in Fig 6A.

## Acknowledgements

Thanks to Dr Teru Kanda for providing B95-8 BAC DNA containing intact FR, and PCR protocols for comparative analyses of FR structure. Thanks to Martin Allday for support and advice and to Paul Farrell for critical reading of the manuscript. Thanks to Peter O’Hare and Thomas Hennig for assistance with microscopy. We thank the St. Mary’s NHLI FACS core facility and their staff, Malte Paulson and Yanping Guo, for support and instrumentation. We thank Beverly Donaldson, Marielle Bouqueau, Thomas Rice and Beate Kampmann for facilitating access to cord and maternal blood samples, collected as part of the MatImms study, and the Imperial College Healthcare NHS Tissue Bank for providing these samples.

